# Liquid-Liquid Phase Separation of the m6A RNA ALKBH9B Demethylase: An IDR-dependent Mechanism driving the Alfalfa Mosaic Virus Infection

**DOI:** 10.64898/2026.07.30.741775

**Authors:** Victor Gonzalez-Silva, Mikhail Oliveira Leastro, Jose Antonio Navarro, Jesús Ángel Sánchez-Navarro, Frederic Aparicio, Vicente Pallás

## Abstract

Biomolecular condensates play crucial role in plant-virus interactions. The plant m6A demethylase ALKBH9B acts as a proviral factor for Alfalfa mosaic virus (AMV), yet the biophysical mechanisms underlying its function remain elusive. Here, we investigate the liquid-liquid phase separation (LLPS) properties of ALKBH9B and their requirements for viral infection. Using *in vitro* and *in vivo* assays (*Nicotiana benthamiana*), we demonstrate that ALKBH9B assembles into highly dynamic, liquid-like cytoplasmic condensates driven primarily by weak hydrophobic interactions. Deletion and mutagenesis analysis identified a C-terminal intrinsically disordered region (IDR3), specifically its arginine-glycine (RG) motifs, as the essential molecular driver of LLPS. Importantly, this domain exhibits dual functionality, mediating both phase separation and direct binding to AMV RNA, which regulates condensate assembly and size. Disruption of the RG motifs completely abolishes ALKBH9B condensation and reduces its proviral capacity, compromising viral accumulation. Collectively, our findings reveal that the spatial compartmentalization of ALKBH9B into dynamic liquid hubs is functionally indispensable for AMV infection. This establishes LLPS as a critical regulatory checkpoint in viral pathogenesis, highlighting how plant viruses exploit the biophysical properties of host epitranscriptomic enzymes to establish specialized, protective microenvironments.

**Highlights:** Our study establishes a direct mechanistic link between the biophysical properties of a plant m6A demethylase and viral pathogenesis, demonstrating that ALKBH9B functions as a key host factor by harnessing LLPS to regulate virus–host interactions.

## INTRODUCTION

Of the approximately 170 internal RNA modifications described to date, the N6-methyladenosine (m6A) is the most abundant internal RNA modification in eukaryotic mRNAs and is dynamically deposited, removed, and recognized by the m6A machinery (Wei and He, 2026). Beyond its essential roles in regulating plant development and responses to abiotic stress (Zhong et al., 2008; Shen et al., 2016; Arribas-Hernández et al., 2018), the m6A machinery has emerged as an important regulator of host-virus interactions (Li et al., 2025; Liu et al., 2026). M6A methylation is catalyzed by a “writer” complex (methyltransferase complex, MTC) while “erasers” remove these marks and YTH-family “readers” modulate the fate of the modified RNAs. By homology with mammals, functional orthologs have been identified in *Arabidopsis*, including the MTC subunits MTA, MTB, FIP37, FIONA, HAKAI, VIRILIZER and HIZ2 (Zhong et al., 2008; Růžička et al., 2017), four ALKBH proteins exhibiting demethylase activity, and *14* YTH-domain readers, comprising the ECT1-11 family and CPSF30 (Arribas-Hernández and Brodersen, 2020).

While MTB and several ECT proteins have been shown to modulate viral RNA stability and fate (Martínez-Pérez et al., 2023; Liu et al., 2025), the role of demethylases is equally decisive. The *Arabidopsis* genome encodes 14 AlkB homologs, of which only ALKBH6, ALKBH9B, ALKBH9C and ALKBH10B have demonstrated *in vivo* m6A demethylase activity; (Duan et al., 2017; Martínez-Pérez et al., 2017; Huong et al., 2020; Amara et al., 2022) whereas indirect evidence suggest that ALKBH8B is a true demethylase (Huong et al., 2022). Notably, ALKBH9B stands out as the only member localized exclusively in the cytoplasm, where it forms granules that colocalize with siRNA bodies and associate with P-bodies (Martínez-Pérez et al., 2017). Besides, analyzing the implication of the protein in the infection cycle of alfalfa mosaic virus (AMV), ALKBH9B was the first demethylase identified to act as a proviral factor in a viral infection (Martínez-Pérez et al., 2017) mediated by its binding to the viral coat protein (CP) and ultimately facilitating viral load in the phloem (Martinez-Perez et al., 2021). Functional mapping of ALKBH9B further revealed that specific structural domains located within its C-terminal intrinsically disordered region (IDR), are required for the physical interaction with both the viral RNA and the AMV CP (Alvarado-Marchena et al., 2021).

AMV is the prototype member of the genus Alfamovirus within the *Bromoviridae* family, whose genome consists of three positive-sense single-stranded RNAs (Thompson et al., 2025). RNA1 and RNA2 encode the replicase complex, while RNA3 is a dicistronic molecule encoding the movement protein (MP) and the CP, the latter being translated from a subgenomic RNA4 (sgRNA4). The CP is a multifunctional protein (Herranz et al., 2012) that, in addition to its role in assembly, is required to initiate the infection through a process known as genome activation, to regulate viral RNA replication and translation and to mediate both cell-to-cell and systemic movement (Bol, 2005; Pallas et al., 2013).

Liquid-liquid phase separation (LLPS) has emerged as a fundamental mechanism for the spatial organization of the cytoplasm through the formation of membraneless organelles, or biomolecular condensates (Brangwynne et al., 2009; Hyman et al., 2014). Interestingly, biomolecular condensates associated with m6A are generated through phase separation and function as dynamic cellular compartments where m6A effectors, RNAs, and regulatory proteins concentrate to orchestrate RNA regulation. Furthermore, m6A itself promotes and modulates condensate assembly (Shen, 2025). These assemblies are typically driven by proteins containing intrinsically disordered regions (IDRs) and low-complexity domains, enriched in disorder-promoting residues, such as glycine (G) and proline (P), as well as arginine (R)-rich motifs, that confer the structural flexibility and multivalency required for the formation of dynamic protein-RNA networks (Gsponer and Madan Babu, 2009; Boeynaems et al., 2017). Such IDR-containing proteins often undergo binding-induced folding and promote the assembly of RNA granules like processing bodies (P-bodies) or stress granules (Lin et al., 2015). In plants, LLPS plays a pivotal role in regulating RNA metabolism and orchestrating stress responses (Zhu et al., 2022; Ostendorp et al., 2022) and emerging evidence indicates that phased transitions contribute substantially to plant-pathogen interactions (Zou et al., 2026). Accordingly, many viruses exploit LLPS to form biomolecular condensates composed of viral and host proteins, thereby concentrating viral components, protecting them from host immune responses, and promoting efficient viral replication (Brown et al., 2021; Fang et al., 2022; Zang et al., 2025; Mei et al., 2026). Among those host proteins, barley m6A demethylase ALKBH1B has recently been shown to undergo phase separation to promote antiviral immunity (Zang et al., 2025). Despite the established importance of the ALKBH9B C-terminal region in binding viral components (Alvarado-Marchena et al., 2021), whether this domain facilitates the assembly of functional condensates through LLPS has not been determined. Although ALKBH9B was previously observed in cytoplasmic granules (Martínez-Pérez et al., 2017), its biophysical nature and the precise molecular requirements underlying its formation remained to be fully characterized.

In this study, we provide a comprehensive biophysical characterization of ALKBH9B phase separation both *in vitro* and *in vivo*. We demonstrate that the C-terminal IDR3 domain and its RG motifs (arginine-glycine-rich region) are necessary to drive the formation of dynamic liquid droplets, which are further modulated by the presence of viral RNA. Importantly, we find that compromising the phase-separating capacity of ALKBH9B correlates with reduced AMV accumulation, suggesting that the spatial organization of this demethylase into biomolecular condensates supports its proviral function. This work links the biophysical state of an m6A ‘eraser’ with its role in viral pathogenesis, providing a new perspective on how viruses exploit host cellular organization to promote infection.

## RESULTS

### The C-terminal IDR3 and its RG motifs are essential drivers of ALKBH9B LLPS

In a previous mutational analysis of ALKBH9B, we identified key regions involved in its function during AMV infection (Alvarado-Marchena et al., 2021). This study predicted the presence of intrinsically disordered regions (IDRs) and mapped the regions responsible for viral CP and RNA binding domain (RBD). Furthermore, those findings led us to propose that i) the C-terminal IDR and the RBD cooperate to promote RNA granules formation and ii) the RGxxxRGG motif located between positions 459-465 plays a critical role in these processes (Alvarado-Marchena et al., 2021).

To investigate the molecular determinants underlying ALKBH9B condensate formation we first mapped the protein domains implicated in this process. **Figure 1A** shows a schematic representation of ALKBH9B, highlighting the central ALKB catalytic domain (orange) flanked by the previously identified CP-binding domain (CP-Bd; yellow) and RBD (blue).

**Figure 1.**
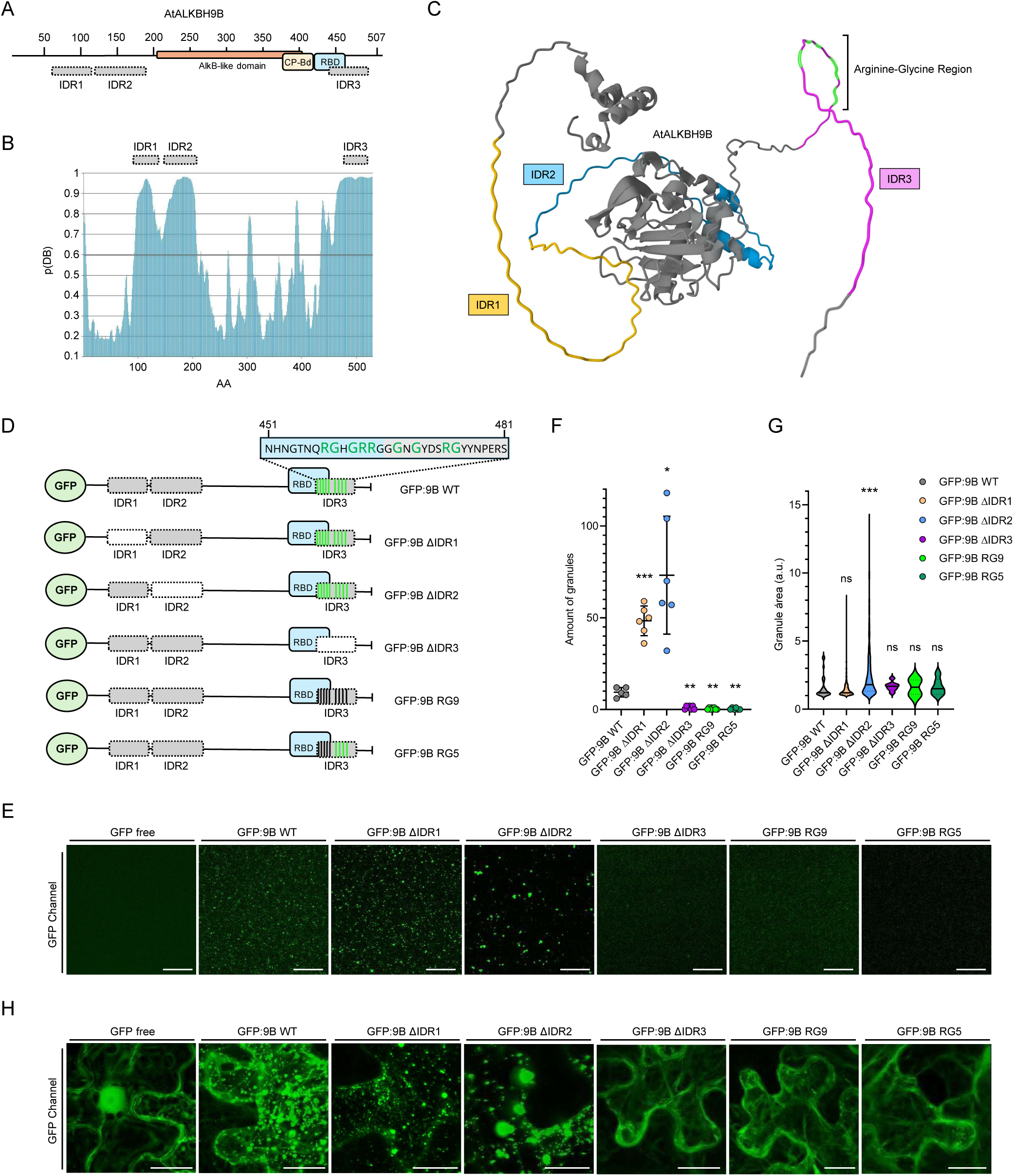
Structural domain architecture of ALKBH9B and functional mapping of its involvement in *in vitro* and *in vivo* condensation. **(A)** Schematic representation of *Arabidopsis thaliana* ALKBH9B (507 aa) highlighting its modular domain architecture: N-terminal intrinsically disordered regions 1 and 2 (IDR1, aa 71-116; IDR2, aa 117-187) and C-terminal IDR3 (aa 457-492) in grey; the central AlkB-like catalytic domain in orange, flanked by the Coat Protein binding domain (CP-Bd) in yellow, and the RNA-binding domain (RBD) in blue. **(B)** Intrinsic disorder propensity profile of ALKBH9B predicted by FuzzDrop, plotting the disorder probability (*p*(DB)) against amino acid position. Horizontal grey bars indicate the selected IDR regions exceeding the structural disorder threshold of 0.6. **(C)** Three-dimensional structural model of ALKBH9B generated by AlphaFold. Structural domains are color-coded as follows: IDR1 in yellow, IDR2 in blue, IDR3 in magenta, arginine glycine (RG) rich cluster highlighted in green. **(D)** Schematic diagram of the N-terminal GFP-fused ALKBH9B construct variants generated for functional mapping, including wild-type (GFP:9B WT), single deletion variants (GFP:9B ΔIDR1, ΔIDR2, ΔIDR3) and alanine-substitution mutants targeting specific arginine and glycine residues within the IDR3 cluster (RG9 and RG5, black lines indicate mutated amino acids). **(E)** Representative *in vitro* confocal microscopy images of purified recombinant free GFP control and GFP:9B variants; scale bars = 20 µm. **(F)** Quantitative analysis of the total number of protein granules formed *in vitro*. Graph displays the mean and ± SEM from six independent images per condition, with individual data points shown. **(G)** Morphometric analysis of the total granular area (expressed in arbitrary units, a.u.) across the different variants *in vitro*. **(H)** Representative *in vivo* confocal microscopy images of *Nicotiana benthamiana* leaf epidermal cells at 3 days post-infiltration, transiently expressing the free GFP control and the indicated GFP:9B construct variants under the control of the constitutive CaMV 35S promoter. The displayed images represent Z-stacks spanning the entire plant cell. Scale bars in (E) and (H) = 20 µm. For quantitative measurements in (F) and (G), statistical significance relative to GFP:9B WT was determined using one-way ANOVA followed by Dunnett’s multiple comparisons test. Asterisks indicate significant differences (ns, non-significant; \**p* < 0.05, \*\**p* < 0.01, \*\*\**p* < 0.001).

To identify regions susceptible to LLPS, we first performed an intrinsic disorder analysis using FuzzDrop (Hatos et al., 2022) (**Fig. 1A and 1B**). The predictions corroborated the presence of three regions with a high probability of disorder (intrinsically disordered regions, IDRs; shown in grey in **Fig. 1A**). Intriguingly, the disorder profile exhibited a pronounced decrease in disorder propensity at the boundary between residues 116 and 117, thereby delineating two independent N-terminal IDRs, designed IDR1 (aa 71-116) and IDR2 (aa 117-187), as well as a C-terminal IDR3 (aa 457-492) overlapping the RNA-binding region. Three-dimensional modeling via AlphaFold supported this organization, showing a structured catalytic core enriched in β-sheets and α-helices, from which the IDRs emerge as highly flexible projections. Within IDR3, specific R and G residues were identified as potential mediators for condensate formation (**Fig. 1C**).

Next, we investigated the contribution of these domains to LLPS both *in vitro* and *in vivo*. To this end, we generate a set of ALKBH9B variants carrying individual deletions of each IDR (mutants ΔIDR1, ΔIDR2, ΔIDR3, respectively) or alanine substitutions within the RG motifs (in green in the box showing sequence in **Fig. 1D**). Two RG mutants were generated: RG9, in which most of the R/G residues in IDR3 were substituted (vertical black lines in IDR3 box in Fig. 1D), and RG5, in which only the residues located within the RBD were mutate (vertical black lines in IDR3 box in Fig. 1D). All proteins were expressed as N-terminal GFP fusions (Fig. 1D). A detailed amino acid sequence alignment comparing the ALKBH9B WT and all these generated mutant variants is provided (**Fig. S1).**

For *in vitro* analysis, *E. coli* BL21 cells transformed with pET28a(+) constructs encoding 9B WT and its mutant variants as N-terminally 6xHis-tagged proteins were used for recombinant protein expression. The proteins were subsequently purified via His-tag affinity chromatography (**Fig. S2**).

We found that GFP:9B WT efficiently formed spherical condensates (**Fig. 1E**). Deletion of either IDR1 or IDR2 did not impair LLPS but instead, significantly increased the number of condensates (*p* < 0.001 and *p* < 0.01, respectively; **Fig. 1F)**, suggesting that these domains negatively regulate condensate nucleation. In contrast, either IDR3 removal or mutation of the RG motifs (RG9 and RG5) completely abolished condensate formation (**Fig. 1E and Fig. 1F**, GFP:9B ΔIDR3, GFP:9B RG9 and GFP:9B RG5, respectively). Interestingly, morphometric analysis showed that GFP:9B ΔIDR2 mutant generated significantly larger granules than the GFP:9B WT (*p* < 0.01; **Fig. 1G**), indicating a specific role for this domain in modulating droplet growth.

Next, we validated these findings *in vivo* through the transient expression of the mutant proteins in *Nicotiana benthamiana* plants. While our *in vitro* assays allowed for a robust quantification of condensate formation (**Fig.1 F, G**), the *in vivo* phenotypes were evaluated qualitatively because they exhibited a clear all-or-none behavior.

Consistent with the *in vitro*results, GFP:9B WT, GFP:9B ΔIDR1 and GFP:9B ΔIDR2 localized to discrete cytoplasmic condensates, whereas the GFP:9B ΔIDR3, GFP:9B RG9 and GFP:9B RG5 mutants exhibited a diffuse cytoplasmic distribution, with no detectable condensate formation (**Fig. 1H, Fig. S3**). Unlike the free GFP control, which is ubiquitously distributed throughout the cell, all ALKBH9B variants were consistently excluded from the nucleus (**Fig. 1H**). Western blot analysis confirmed protein expression and stability (**Fig. S4**). The condensate-forming capacity of GFP:9B WT was maintained following transient expression under the control of its native promoter, precluding the possibility that granule formation resulted from overexpression artifacts (**Fig. S5**). Taken together, these results demonstrate that IDR3 and its RG motifs are essential for ALKBH9B condensate formation.

### ALKBH9B undergoes *in vitro* aggregation driven by hydrophobic forces

The physical state of proteins undergoing phase separation is typically governed by protein concentration and the surrounding molecular environment (Hyman et al., 2014; Boeynaems et al., 2018). To characterize these properties, we analyzed the behavior of purified GFP:9B WT subjected to different conditions by confocal microscopy assays.

The initial experiments were focused on the impact of protein concentrations ranging from 1 to 24 µM. WT ALKBH9B exhibited a dose-dependent propensity for biocondensate formation, showing a progressive increase in spherical condensates as protein levels increased (**Fig. 2A**). In contrast, free GFP remained diffusely distributed across the entire concentration range examined, without forming biomolecular condensates (**Fig. 2A**). Quantification revealed a positive correlation between GFP:9B WT concentration and total condensate area, reaching statistical significance at 24 µM (*p* < 0.05; **Fig. 2D**). These data point to the existence of a critical concentration required for the nucleation of stable ALKBH9B condensates.

**Figure 2.**
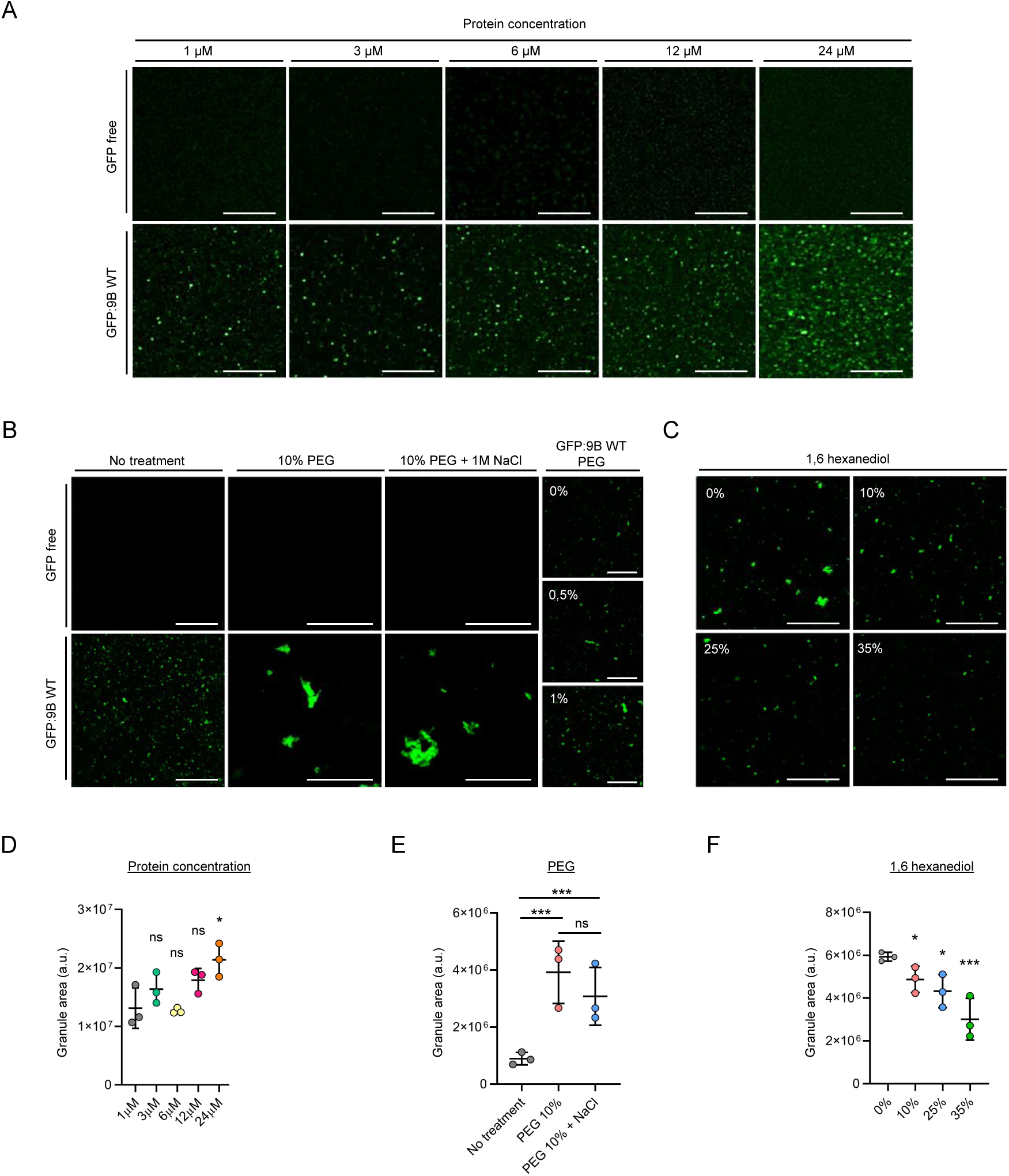
*In vitro* macromolecular condensation of ALKBH9B under varying protein concentrations, crowding agents, and chemical disruptors. **(A)** Representative *in vitro* confocal microscopy images of purified recombinant free GFP control and GFP:9B WT at increasing protein concentrations (1, 3, 6, 12 and 24 µM). **(B)** Representative *in vitro* confocal microscopy images showing the effect of macromolecular crowding (PEG 8000) and ionic strength on free GFP and GFP:9B WT. Panels include a concentration gradient of PEG (0%, 0.5% and 1%), an untreated control, a 10% PEG treatment, and a combination of 10% PEG with 1 M NaCl. **(C)** Representative *in vitro* confocal microscopy images of GFP:9B WT granules treated with increasing concentrations of 1,6-hexanediol (0%, 10%, 25% and 35%). **(D)** Morphometric quantification of total granular area (expressed in arbitrary units, a.u.) across the indicated GFP:9B WT protein concentrations. **(E)** Morphometric quantification of total granular area (a.u.) under untreated, 10% PEG, and 10% PEG + 1M NaCl conditions. **(F)** Morphometric quantification of total granular area (a.u.) following treatment with the indicated concentration of 1,6-hexanediol. Scale bars in (A-C) = 12.5 µm. For all quantitative data (D-F), graphs display the mean ± SEM from three independent images per condition, with individual data points shown. Statistical significance was determined using one-way ANOVA followed by Dunnett’s multiple comparisons test relative to the respective controls in (D) (1 µM) and (F) (0%), or by Tukey’s multiple comparisons test for pairwise comparisons in (E) (ns, non-significant; \**p* < 0.05, \*\*\**p* < 0.001).

To mimic the crowded intracellular environment, we evaluated the effect of polyethylene glycol (PEG 8000) as a macromolecular crowding agent. The addition of 10% PEG enhanced GFP:9B WT LLPS, yielding larger and more numerous condensates (**Fig. 2B**). This promotion of phase separation followed a dose-dependent pattern, with visible granulation even at PEG concentrations as low as 0.5% and 1% (**Fig. 2B**). Inclusion of 10% PEG resulted in a substantial increase in condensate area compared to the untreated protein (*p* < 0.001; **Fig 2E**).

The molecular forces stabilizing these condensates were probed by assessing their sensitivity to ionic strength and chemical disruptors. The addition of 1 M NaCl failed to alter the stability of PEG-induced condensates, and no significant differences were observed between 10% PEG alone and the PEG-NaCl combination (**Fig. 2B, E**). This observation suggests that electrostatic interactions are not the primary drivers of condensate cohesion. In contrast, treatment with the aliphatic alcohol 1,6-hexanediol, a compound known to disrupt many biomolecular condensates when present at sufficient concentrations by breaking weak hydrophobic interactions (Kroschwald et al., 2017; Zhu et al., 2022), dissolved GFP:9B WT granules in a dose-dependent manner (**Fig. 2C**). The reduction in granular area reached significance at 10% (*p* < 0.05) and was most pronounced at 35% (*p* < 0.001; **Fig. 2F**). Altogether, these findings demonstrate that ALKBH9B undergoes *in vitro* LLPS driven primarily by hydrophobic interactions and facilitated by macromolecular crowding.

### ALKBH9B undergoes *in vivo* phase separation driven by hydrophobic forces

To determine whether the biophysical properties observed *in vitro* extend to a cellular context, we analyzed ALKBH9B localization via transient expression in *N. benthamiana* leaves. Microscopy confocal imaging confirmed that GFP:9B WT predominantly localizes to the cytoplasm as discrete condensates (**Fig. 1H**). Given that the pronounced effects of the chemical treatments on condensate formation and dissolution were visually distinguishable by microscopy, the *in vivo* dynamics were evaluated qualitatively.

The impact of macromolecular crowding was evaluated by infiltrating increasing concentrations of PEG 8000. In plants expressing GFP:9B WT, PEG induced a dose-dependent increase in cytoplasmic condensate formation; with condensates increasing in both number and size and often clustering in perinuclear regions while remaining strictly excluded from the nucleus (**Fig. 3A**). This promoting effect persisted upon the addition of NaCl 1 M, supporting the observation that condensate stability is not primarily mediated by electrostatic forces (**Fig. 3A**). In contrast, the free GFP control remained diffusely distributed throughout the cytoplasm and nucleus without forming granules. Meanwhile, the RG5 mutant failed to form condensates under any of the tested osmotic stress conditions, exhibiting a diffuse cytoplasmic signal even at 10% PEG (**Fig. 3A**). These findings establish that RG motifs are essential molecular requirements for ALKBH9B LLPS, even under high-crowding conditions.

**Figure 3.**
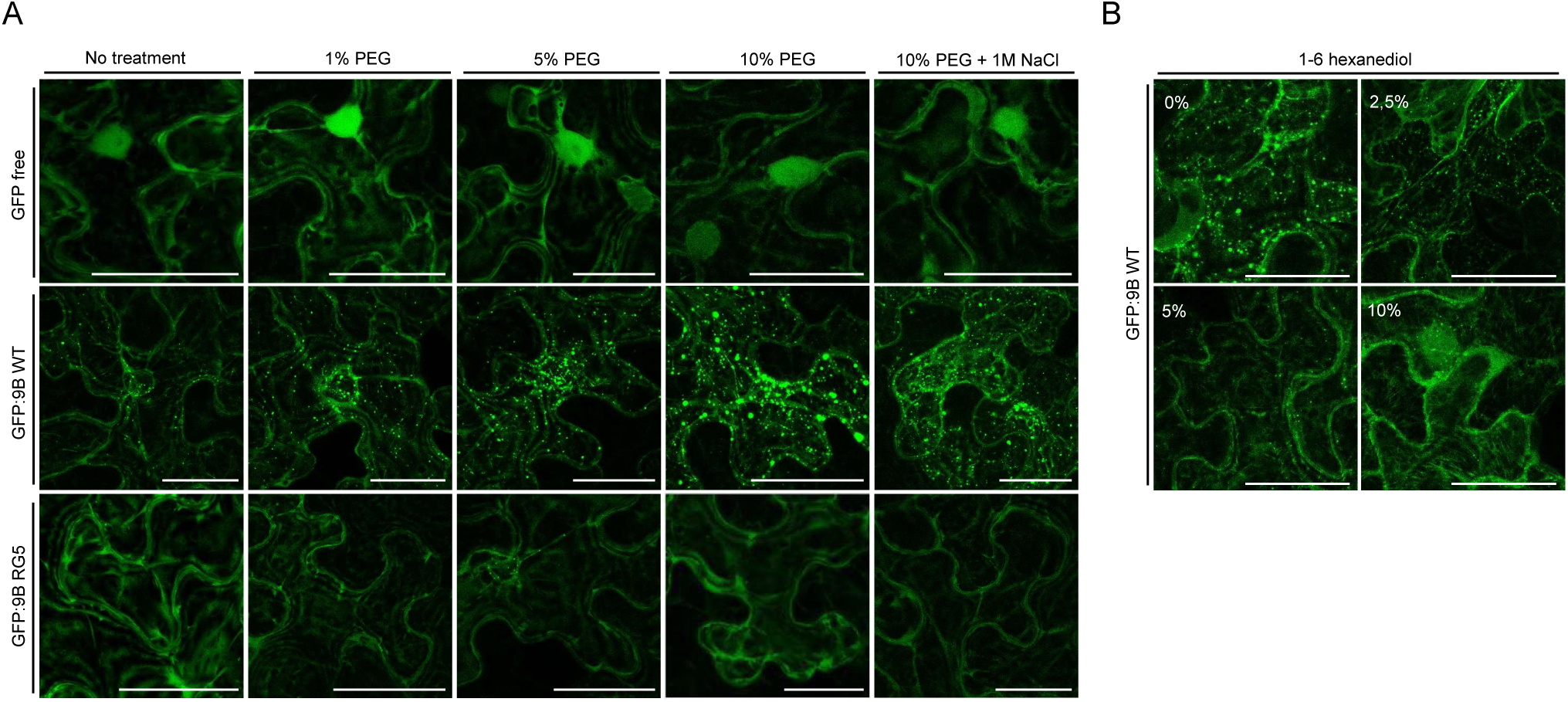
*In vivo* condensation of ALKBH9B under macromolecular crowding, ionic strength, and chemical disruptors. **(A)** Representative *in vivo* confocal microscopy images of *Nicotiana benthamiana* leaf epidermal cells at 2 days post-infiltration, transiently expressing the free GFP control, GFP:9B WT, and the GFP:9B RG5 mutant. The panels show cells subjected to an untreated control condition, a concentration gradient of macromolecular crowding agent (1%, 5% and 10% PEG 8000), and a combination of 10% PEG with 1 M NaCl. **(B)** Representative *in vivo* confocal microscopy images of *Nicotiana benthamiana* leaf epidermal cells transiently expressing GFP:9B WT following treatment with increasing concentrations of 1,6-hexanediol (0%, 2.5%, 5%, and 10%). The displayed images represent Z-stacks spanning the entire plant cell volume. Scale bars in (A) and (B) = 20 µm.

We next examined the sensitivity of ALKBH9B granules to 1,6-hexanediol. The treatment induced a dose-dependent dissolution of the ALKBH9B condensates (**Fig. 3B**). At 10% hexanediol, the GFP:9B WT signal became diffuse and partially relocalized to tubular structures resembling the cytoskeleton, with occasional accumulation within the nuclear compartment (**Fig. 3B**). This delocalization suggests that high hexanediol concentrations may compromise organellar membrane integrity or cytoskeletal organization, thereby disrupting protein homeostasis. Collectively, these data confirm that *in vivo* ALKBH9B condensation is a dynamic process driven by hydrophobic interactions and mediated by RG motifs.

### Intracellular environment dictates the liquid properties and high internal mobility of ALKBH9B biocondensates

To characterize ALKBH9B biochemical properties and molecular dynamics, we performed fluorescence recovery after photobleaching (FRAP) assays under both *in vitro* and *in vivo* conditions.

Under *in vitro* conditions, GFP:9B WT granules exhibited highly restricted internal mobility. Following photobleaching in the absence of PEG, no detectable fluorescence recovery was observed throughout period (**Fig. 4A**). Kinetic analysis confirmed this immobile state, with fluorescence remaining near post-bleach levels (∼14%) and precluding reliable estimation of the half time (t_1/2_) of recovery because of the absence of measurable molecular exchange (**Fig. 4B**). Increasing the concentration of the crowding agent to 20% PEG yielded similar results; despite a slight increase in the fluorescence plateau (∼22%), recovery remained minimal (**Fig. 4C**), suggesting that isolated ALKBH9B forms highly viscous or gel-like condensates under these conditions. In contrast to the *in vitro* findings, ALKBH9B condensates displayed dynamic behavior and liquid-like properties within the cellular environment. Time-lapse imaging revealed granule fusion events, in which adjacent cytoplasmic condensates approached each other and merged into larger structures (**Fig. 4D**). Fusion kinetics varied considerably, ranging from slower events requiring up to ∼40 seconds (**Fig. 4D**, upper panel) to rapid coalescence (∼2 seconds, between second 12 and 14, **Fig. 4D** lower panel). The fluid nature of these *in vivo* condensates was validated by FRAP. Unlike the *in vitro* results, cytoplasmic GFP:9B WT granules showed rapid and substantial fluorescence recovery after bleaching (**Fig. 4E**). One-phase association curve fitting revealed a mobile fraction of 65.8% (Plateau = 0.66), indicating that most ALKBH9B molecules within the condensate undergo constant exchange with the surrounding cytoplasm (**Fig. 4F**). Recovery kinetics were remarkably rapid, with a half-time (t_1/2_) of 4.06 seconds (**Fig. 4F**). These results demonstrate that while ALKBH9B possesses an intrinsic tendency for stable aggregation, the cellular environment maintains these condensates in a highly dynamic liquid state, facilitating molecular exchange.

**Figure 4.**
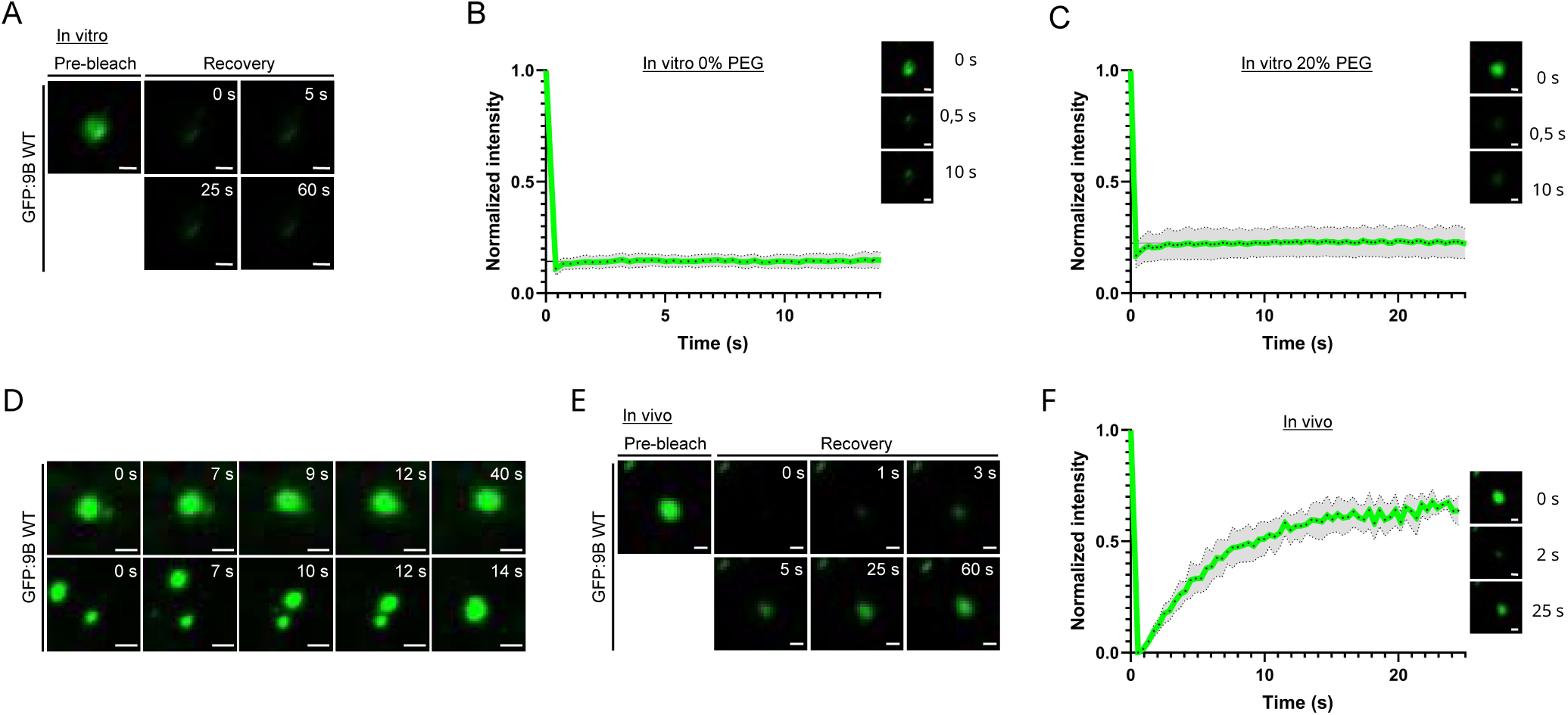
*In vitro* and *in vivo* molecular dynamics and liquid-like properties of ALKBH9B condensates. **(A)** Representative *in vitro* fluorescence recovery after photobleaching (FRAP) time-lapse images of purified GFP:9B WT granules in the absence of PEG. Images show the pre-bleach state and recovery at 0, 5, 25, and 60 s post-bleaching. **(B)** Normalized *in vitro* FRAP recovery curve for GFP:9B WT granules (0% PEG). Kinetic analysis indicates a mobile fraction plateau of ∼14%. Inset images show the targeted granule at pre-bleach (0 s), 0.5 s and 10 s post-bleaching. **(C)** Normalized *in vitro* FRAP recovery curve for GFP:9B WT granules in the presence of 20% PEG. Kinetic analysis indicates a mobile fraction plateau of ∼22%. Inset images display the targeted granule at pre-bleach (0 s), 0.5 s, and 10 s post-bleaching. **(D)** Representative *in vivo* time-lapse images showing granule fusion events of GFP:9B WT condensates in *Nicotiana benthamiana* leaf epidermal cells. The top panel sequence captures a slower fusion event (0, 7, 9, 12, and 40 s), while the bottom panel sequence illustrates a rapid fusion event (0, 7, 10, 12, and 14 s). **(E)** Representative *in vivo* FRAP time-lapse images of cytoplasmic GFP:9B WT condensates, displaying the pre-bleach state and recovery at 0, 1, 3, 5, 25 and 60 s post-bleaching. **(F)** Normalized *in vivo* FRAP recovery curve for GFP:9B WT condensates. Kinetic parameters obtained from a one-phase association curve fitting reveal a mobile fraction of 65.8% and a recovery half-time (t_1/2_) of 4.06 s (inset images display the targeted condensate at pre-bleach (0 s), 2 s and 25 s post-bleaching). For all FRAP recovery graphs (B, C, and F), the green dotted line represents the mean normalized intensity of 5 independent FRAP experiments (performed on individual distinct granules), and the grey shaded area indicates the standard deviation. Scale bars in (A-F) = 1 µm.

### IDR3 and its RG motifs mediate RNA interaction and modulate condensate dynamics

Given the overlap between IDR3 and the predicted RBD, we investigated whether ALKBH9B phase separation is linked to its RNA-binding function. We performed a northwestern blot assay using an *in vitro* transcript of AMV sgRNA4. Coomassie Brilliant Blue staining confirmed uniform loading and integrity for all purified protein variants (**Fig. 5A**, upper panel). Northwestern assays revealed robust RNA-binding for GFP:9B WT, GFP:9B ΔIDR1 and GFP:9B ΔIDR2 (**Fig. 5A**, lower panel). In contrast, deletion of the C-terminal domain (GFP:9B ΔIDR3) or mutation of the RG motifs (GFP:9B RG9 and GFP:9B RG5) completely abolished RNA-binding capacity, yielding a lack of signal similar to the free GFP control (**Fig. 5A**, lower panel). This data reinforce and extend our previous proposal (Alvarado-Marchena et al., 2021), indicating that IDR3, and specifically its RG motifs, constitute the molecular determinants required for RNA recognition by ALKBH9B.

**Figure 5.**
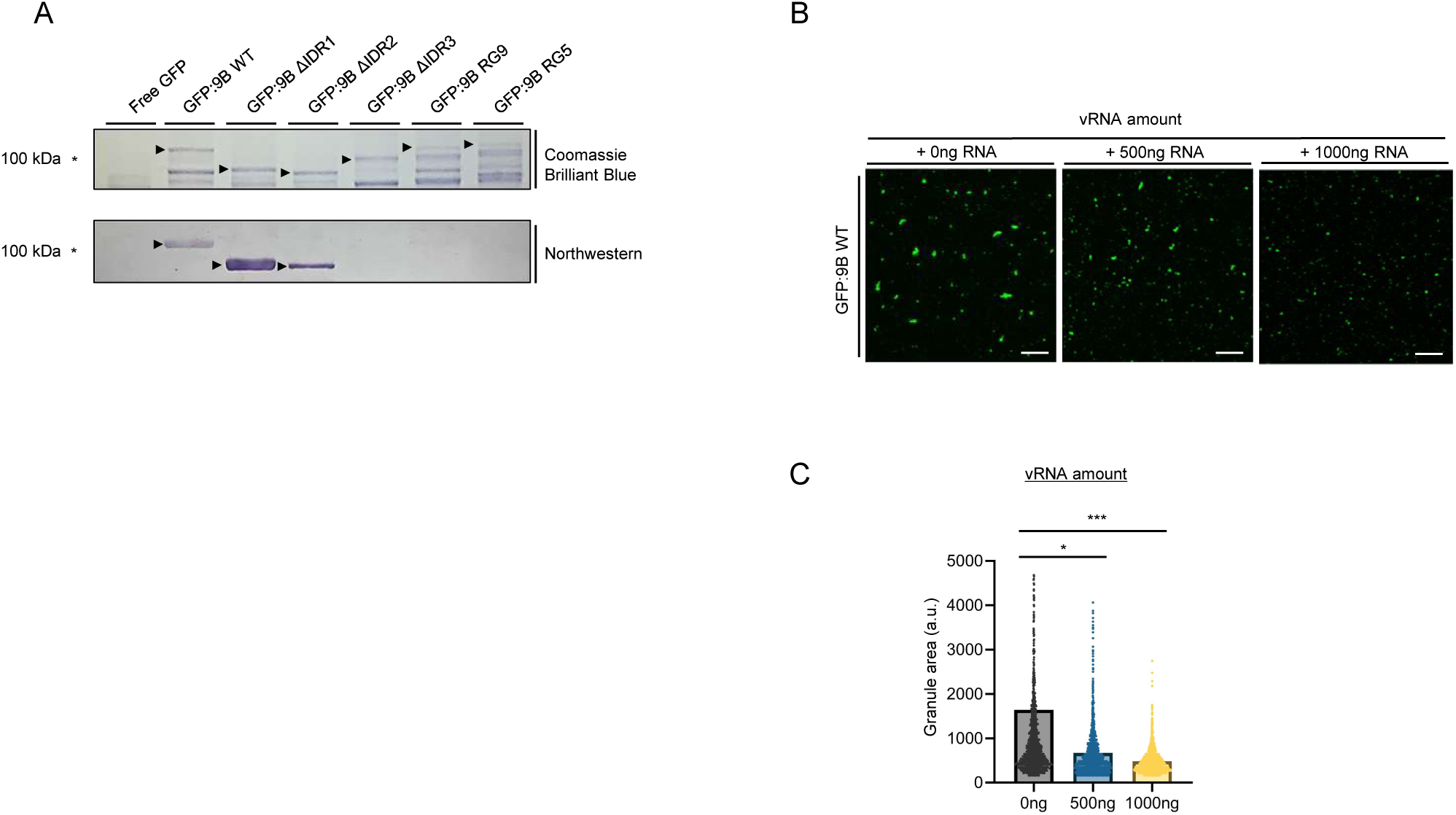
RNA-binding capacity of ALKBH9B variants and RNA-dependent modulation of *in vitro* condensation. **(A)** Northwestern blot analysis of the interaction between purified GFP:9B variants and viral RNA. Free GFP was included as a negative control. The upper panel shows Coomassie Brilliant Blue staining of the recombinant proteins to confirm equal loading. The lower panel shows the Northwestern blot probed with a DIG-labeled *in vitro* transcript of AMV sgRNA4. Arrowheads indicate the expected position of the corresponding protein bands. **(B)** Representative *in vitro* confocal microscopy images of GFP:9B granules incubated with increasing amounts of the AMV sgRNA4 transcript (0, 500, and 1000 ng). **(C)** Morphometric quantification of total granular area (expressed in arbitrary units, a.u.) across the indicated RNA amounts. Scale bar in (B) = 10 µm. For the quantitative data in (C), graphs display the mean ± SEM with individual data points shown. Statistical significance relative to the 0 ng RNA control was determined using one-way ANOVA followed by Dunnett’s multiple comparisons test (\**p* < 0.05, \*\*\**p* < 0.001).

After establishing this binding capacity, we examined whether the presence of RNA modulates the physical properties of ALKBH9B condensates *in vitro*. Confocal microscopy showed that the addition of increasing viral RNA amounts (0, 500, and 1000 ng) altered the size of GFP:9B WT granules (**Fig. 5B**). As RNA availability in the mixture increased, the condensates became progressively smaller, suggesting an RNA-mediated molecular dispersion or reorganization effect (**Fig. 5B**).

Quantification of the granular area supported these visual observations. The addition of 500 ng of RNA resulted in a significant reduction in condensate size (*p* < 0.05). This effect was more pronounced at 1000 ng, where the granular area decreased markedly (*p* < 0.001) compared to the RNA-free condition (**Fig. 5C**). Together, these results indicate that IDR3 is a bifunctional domain indispensable for both RNA binding and phase separation, with RNA acting as a critical regulator of ALKBH9B condensate size and abundance.

### ALKBH9B phase separation and RNA interaction determine its proviral function

ALKBH9B is a known proviral factor facilitating AMV infection (Martínez-Pérez et al., 2017). To determine whether this role depends on biomolecular condensate formation and RNA binding, we assessed AMV accumulation using phase-separation-deficient ALKBH9B variants by overexpression analysis. We performed agroinfiltration assays in *N. benthamiana* leaves, co-expressing AMV genomic RNAs with GFP:9B WT, GFP:9B ΔIDR3, GFP:9B RG5, or an empty vector (pMOG800) as control. Viral accumulation was analyzed three days post-infiltration via northern blot to detect viral RNAs with signal intensities normalized to rRNA loading. A representative blot from the three independent bioassays is shown (**Fig. 6A**).

**Figure 6.**
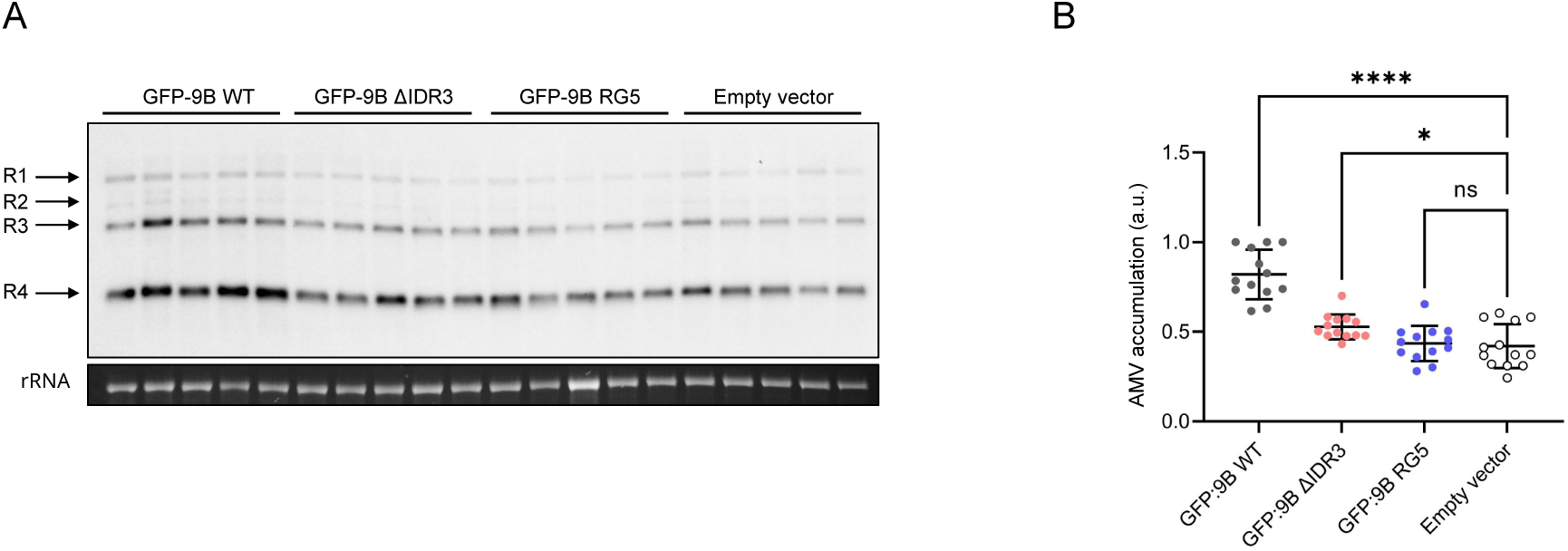
Effect of ALKBH9B variants on Alfalfa mosaic virus (AMV) accumulation *in vivo*. **(A)** Representative Northern blot analysis of AMV RNA accumulation in *Nicotiana benthamiana* leaves at 3 days post-infiltration. Leaves were co-infiltrated with AMV genomic RNAs and either an empty vector (pMOG) control, GFP:9B WT, GFP:9B ΔIDR3, or the GFP:9B RG5 mutant. The upper panel displays the viral RNAs detected using a mixture of probes specific for the three AMV genomic RNAs and the subgenomic RNA 4. The lower panel shows the rRNA bands used as a loading control. The Northern blot shown is representative of three independent bioassays. **(B)** Quantification of total viral RNA accumulation normalized to rRNA levels across the indicated conditions. For the quantitative data in (B), the graph displays the mean ± SEM of pooled data from the three independent biological replicates (*n* = 13 total samples per condition), with individual data points shown. Statistical significance relative to the empty vector control was determined using one-way ANOVA followed by Dunnett’s multiple comparisons test (ns, non-significant; \**p* < 0.05, \*\*\*\**p* < 0.0001).

Northern blot quantification pooling all data points across the three independent biological replicates revealed that GFP:9B WT overexpression led to significantly higher viral accumulation compared to the empty vector control (*p* < 0.0001, **Fig. 6B**), confirming its proviral role. In contrast, the mutant unable to form granules due to substitution of its arginine-glycine residues, GFP:9B RG5, showed no statistically significant differences in viral accumulation relative to the empty vector control (**Fig. 6B**), indicating a complete loss of its proviral capacity. Interestingly, the deletion variant GFP:9B ΔIDR3 retained a partial, intermediate effect, exhibiting a modest but significant increase in viral levels compared to the empty vector control (*p* < 0.05, **Fig. 6B**), although this significance was only supported by one of the three individual bioassays (**Fig. S6**).

Taken together, these data demonstrate that the proviral function of ALKBH9B is strictly linked to its C-terminal IDR3. Our findings suggest that cytoplasmic condensate formation and/or RG-mediated RNA binding are essential requirements for ALKB9B to promote AMV accumulation in plants.

## DISCUSSION

Liquid–liquid phase separation (LLPS) has recently emerged as a fundamental mechanism for organizing cellular processes and as a key regulatory hub in plant–virus interactions (Mei et al., 2026). Likewise, epitranscriptomic regulation through LLPS has been recognized as a major mechanism coordinating RNA metabolism during plant development and stress responses (Shen, 2025). Notably, several components of the m6A RNA modification machinery have been shown to undergo LLPS (Wu et al., 2024; Zang et al., 2025), suggesting that condensate formation is an integral feature of m6A-mediated RNA regulation. However, the molecular mechanisms governing the assembly of these condensates and their contribution to the plant viral infection cycle remain largely unknown. Intrinsically disordered regions (IDRs), which drive multivalent interactions, are essential for the assembly and function of most biomolecular condensates. The Arabidopsis m6A RNA demethylase ALKBH9B (Martínez-Pérez et al., 2017) contains three predicted IDRs located at the N-terminal (IDR1 and IDR2), and C-terminal (IDR3) regions and an RNA-binding domain (RBD) within the latter (Alvarado-Marchena et al., 2021), making it an attractive model to investigate the role of phase separation in antiviral RNA regulation. An increasing body of evidence supports the notion that phase-separating RNA-binding proteins (RBPs) containing RBDs and IDRs are critical for plant development and adaptation to environmental stress (Fan et al., 2024). Here, we demonstrate that the C-terminal arginine-glycine rich motif (RG5) located at the IDR3 is the primary driver of ALKBH9B LLPS. Arginine-rich intrinsically disordered regions are well known to facilitate multivalent biomolecular interactions via cation-π and stacking forces (Li et al., 2012; Boeynaems et al., 2017).

Our chemical perturbation assays refined this view by revealing that, although arginine residues are commonly associated with electrostatic interactions, the resistance of ALKBH9B condensates to high NaCl concentrations, together with their rapid dissolution upon 1,6-hexanediol treatment, indicates that weak hydrophobic interactions within the RG5 cluster dominate the condensation process, revealing a high internal dynamism.

Furthermore, our data demonstrates that the RG5 motif also mediates viral RNA binding (Alvarado-Marchena et al., 2021), highlighting the dual functional role of this region. *In vitro*, viral RNA modulates ALKBH9B condensation organization in a concentration-dependent manner. Rather than acting as a passive component, the viral genome may function as a nucleating scaffold at appropriate protein-to-RNA ratios while limiting condensate growth at higher RNA concentrations, a behavior consistent with the reentrant phase separation described for several RNA-biding proteins (Maharana et al., 2018). A similar situation has been observed for the *Arabidopsis* phloem-associated RNA chaperone-like (PARCL) protein, in which RNA binding and condensate formation can be independently regulated through the C-terminal region and/or the IDR (Ostendorp et al., 2022). This observation suggests that ALKBH9B condensation is strictly regulated by the stoichiometry relationship between viral RNA and the demethylase, potentially coupling condensate assembly to the progression of viral replication.

In a previous work, we reported that ALKBH9B co-localizes with markers of different cytoplasmic granules, such as SUPPRESSOR OF GENE SILENCING 3 (SGS3), mRNA-decapping enzyme 2 (DCP2) and upstream frameshift 1 (UPF1) components of siRNA bodies, P-bodies and nonsense-mediated mRNA decay system (NMD), respectively (Martinez-Pérez et al., 2017), although the material and biophysical properties of these assemblies remained unknown. Interestingly, it has been recently demonstrated that under heat stress conditions, the m6A demethylase ALKBH9B localizes to stress granules (SGs) along with m6A-modified transcripts where selectively demethylates the heat activated retroelement Onsen, leading to the subsequent release of Onsen from SGs (Fan et al., 2023). The present work clarifies that these cellular granules are not static or rigid aggregates, but highly dynamic, liquid-like condensates. *In vivo* transient expression in *N. benthamiana* provided definitive proof of this fluid state by revealing spontaneous droplet fusion events. This liquid nature was further quantified by FRAP analysis, showing a substantial mobile fraction of 65% and rapid fluorescence recovery. This high molecular exchange is functionally critical; the internal dynamism maintained by weak hydrophobic interactions allows the ALKBH9B enzyme to rapidly diffuse, leading to its compartmentalization into biomolecular condensates, which creates not only an optimized enzymatic microenvironment to enhance its demethylase efficiency but could also facilitate the coordinated, multi-component assembly of these biocondensates.

A striking evolutionary contrast exists between our findings and the recently characterized barley demethylase, HvALKBH1B (Zang et al., 2025). While both erasers undergo LLPS to regulate RNA virus infections, their physiological outputs are opposed. HvALKBH1B condenses to orchestrate host antiviral restriction against rhabdoviruses, whereas ALKBH9B phase separation functions as a strictly proviral mechanism supporting AMV accumulation. This discrepancy demonstrates that the biological outcome of epitranscriptomic phase separation is highly context-dependent, likely governed by the specific macromolecular composition of the condensate (Wang et al., 2018). While antiviral aggregates typically sequester and isolate viral replication machinery to suppress infection (Fang et al., 2022), ALKBH9B exploits its dynamic liquid state to establish a specialized microenvironment. In this niche, the enzyme outcompetes host defenses by selectively processing the co-condensed viral genome. Consequently, these data highlight how plant viruses can subvert or repurpose conserved biophysical pathways to convert a potential defense mechanism into a susceptibility factor (Brown et al., 2021).

Our results further demonstrate that disrupting ALKBH9B phase separation through mutation of the RG5 motif significantly reduces AMV accumulation at early times of infection, establishing a direct link between spatial condensation and proviral activity. Previous work from our laboratory established a model wherein ALKBH9B-mediated m6A removal maintains the AMV infection in a hypomethylated state, thereby evading the host counter-defense mechanism driven by the YTH-domain readers ECT2, ECT3, and ECT5 (Martínez-Pérez et al., 2023).

Our current results expand this model by revealing that this viral shielding is spatially regulated: the compartmentalization of ALKBH9B within dynamic liquid hubs is required for optimal m6A removal. Conversely, when phase separation is compromised, the viral RNA remains hypermethylated and ECT reader effectors recognize the persistent m6A marks and, likely driven by their own IDRs, effectively prevent its access to the translation machinery and virion assembly (**Fig. 7**). In this context, the inability of the ALKBH9B-RG5 mutant to associate with condensates may allow hypermethylated viral RNAs to be occupied by multiple ECT proteins, potentially driving their sequestration and subsequent destabilization. We cannot discard the possibility that specific host m6A mRNAs required for efficient viral infection would be affected in the same way. Consequently, the biophysical function of ALKBH9B to form functional condensates represents, in addition to the ECT axis, a critical regulatory checkpoint in plant-virus co-evolution, enabling the virus to evade degradation pathways.

**Figure 7.**
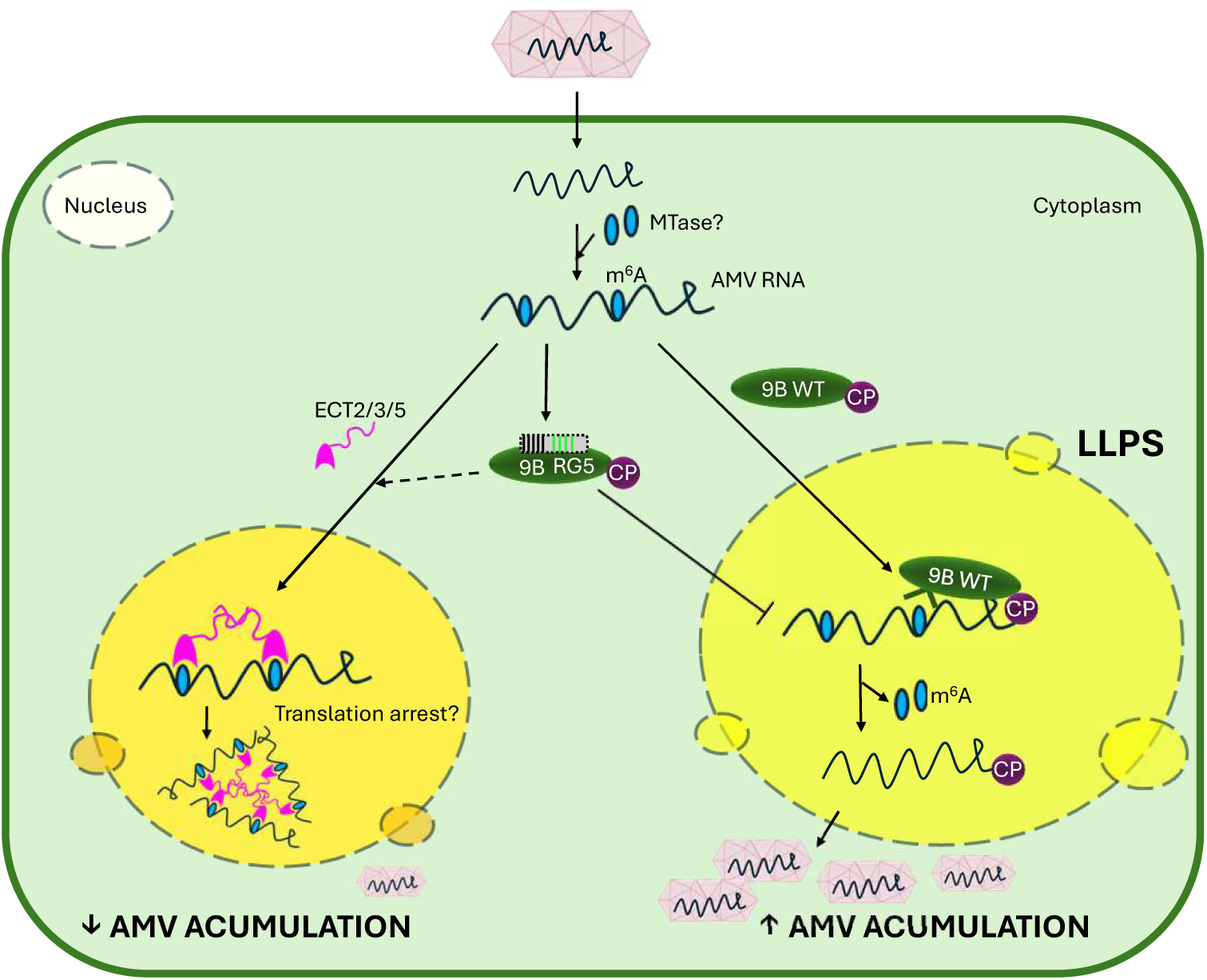
Proposed working model for the proviral function of ALKBH9B during Alfalfa mosaic virus (AMV) infection. Schematic representation of the interplay between AMV RNA, host proteins and phase separation dynamics. Upon infection, AMV RNA is methylated by a host methyltransferase (MTase). In a functional context, wild-type ALKBH9B recognizes and interacts with the methylated viral RNA via its C-terminal RG motifs. This interaction drives the recruitment of the viral transcripts into cytoplasmic liquid-liquid phase separation (LLPS) condensates. Within these granules, ALKBH9B mediates viral RNA demethylation. While ALKBH9B is also known to interact with the viral coat protein (CP), it remains an open question whether this association takes place within the condensates or in the surrounding cytoplasm. Ultimately, this demethylation step prevents host surveillance and maintains the viral RNA in a state that is highly accessible to the translation and replication machinery, thereby promoting successful viral accumulation. Conversely, when ALKBH9B lacks a functional RNA-binding domain (such as the RG5 mutant), it fails to interact with the AMV RNA and does not localize into cytoplasmic granules, suggesting that RNA binding and condensate formation are closely coupled processes. As a result, the methylated viral RNA is left unprotected and becomes recognized by host reader proteins, such as ECT2/3/5. These host factors interact with the viral RNA and, driven by their own intrinsically disordered regions (IDRs), sequester the viral transcripts away from the replication and translation pools, ultimately restricting viral accumulation. In the diagram, ALKBH9B (9B), the viral CP and host ECT proteins are depicted in green, purple, and pink, respectively. Small blue circles (m6A) denote RNA methylation marks added by an unidentified methyltransferase (MTase ?), and the yellow shaded area represents the LLPS condensate within the cytoplasm.

In conclusion, our work establishes a mechanistic link between the biophysical properties of a plant m6A demethylase and viral pathogenesis, demonstrating that ALKBH9B functions as a key host factor by harnessing LLPS to regulate virus-host interactions (Brown et al., 2021). Future studies should determine whether the condensate environment intrinsically enhances demethylase kinetics, and how the precise *in vivo* stoichiometry between viral RNA, the AMV CP, and the host enzyme dictates the assembly or dissolution of these dynamic viral hubs. Ultimately, decoding these macromolecular transitions opens promising, broad-spectrum bioengineering avenues to protect crops by strategically disrupting the biophysical condensation mechanism that plant viruses exploit to thrive.

## Materials and Methods

### Plant material and growth conditions

Wild-type *N. benthamiana* plants were grown in a controlled-environment chamber under standard conditions. The photoperiod was maintained at 16 hours of light and 8 hours of dark, with a temperature of 24°C-22°C and a relative humidity of approximately 60%. Plants were used for agroinfiltration assays at 4 to 5 weeks of age.

### Bioinformatic analysis and structural prediction

The identification of potential LLPS domains and Intrinsically Disordered Regions (IDRs) within the ALKBH9B sequence was performed using the FuzzDrop web server (Hatos et al., 2022). The three-dimensional (3D) structure of the ALKBH9B protein was predicted using AlphaFold (Jumper et al., 2021). The resulting structural models and the spatial distribution of the predicted IDRs were visualized and analyzed using PyMOL software. Additionally, amino acid sequence alignments of ALKBH9B WT and its mutant variants were performed using Clustal Omega and visualized with ESPript 3.2.

### Plasmid construction and mutagenesis

For *in planta* expression assays, the coding sequence of ALKBH9B WT and its mutant variants were fused in frame to the GFP. The specific oligonucleotides used for the construction of the original plasmid and the generation of all deletion and substitution mutants are listed (**Table S1**). These fusion cassettes were cloned into the pMOG binary vector. Depending on the assay, transcription was driven by either the constitutive Cauliflower mosaic virus (CaMV) 35S promoter or the endogenous ALKBH9B native promoter, and terminated by the Popit terminator. All constructs were first transformed into *Escherichia coli* DH5α competent cells for plasmid amplification and sequence verification. Verified binary plasmids were subsequently transformed into *Agrobacterium tumefaciens* strains C58 by electroporation for transient expression assays.

### Agrobacterium-mediated transient expression and viral infection

For transient expression and viral infection assays, *Agrobacterium tumefaciens* C58 strain carrying the respective constructs were grown overnight at 28°C. Cells were harvested by centrifugation and resuspended in infiltration buffer (10 mM MES pH 5.6, 10 mM MgCl_2_). To evaluate viral accumulation and conduct *in vivo* Fluorescence Recovery After Photobleaching (FRAP) experiment, *N. benthamiana* leaves were co-infiltrated with a specific mixture of *Agrobacterium* cultures. The *AMV* infectious clone components (RNA1, RNA2 and RNA3) were infiltrated at a final optical density (*OD_600_*) of 0.1 for each, whereas GFP:9B WT and its mutant variants (ΔIDR1, ΔIDR2, ΔIDR3, RG5, and RG9) were infiltrated at a final *OD_600_* of 0.5. Depending on the specific assay, 9B constructs were expressed either under the control of the constitutive CaMV 35S promoter or its endogenous native promoter, as indicated in the text. The bacterial suspensions were infiltrated into the abaxial side of 4-to 5-week-old *N. benthamiana* leaves using a needleless syringe.

### RNA extraction and Northern blot analysis

Viral RNA accumulation was evaluated by Northern blot analysis. Infected leaf tissues were harvested at 4 days post-agroinfiltration and ground to a fine powder in liquid nitrogen using a mortar and a pestle. Total RNA was extracted from 100 mg of homogenized leaf material using TRIzol Reagent (Invitrogen). Following phase separation with chloroform, the aqueous phase was recovered, and RNA was precipitated with isopropanol. The resulting RNA pellet was further purified and concentrated by an additional precipitation step using 3 M sodium acetate and absolute ethanol. For each sample, 500 ng of purified total RNA was denatured by formaldehyde treatment, resolved by electrophoresis on a 1.2% formaldehyde-agarose gel, and capillary-transferred to a positively charged nylon membrane (Roche). Viral transcripts were detected using specific digoxigenin (DIG)-labeled riboprobes complementary to the four viral RNAs of *AMV*. The synthesis of DIG-labeled riboprobes, membrane hybridization conditions, and subsequent DIG-detection procedures were performed as previously described (Pallás et al., 1998).

### 9B expression and purification

The 9B WT, 9B ΔIDR1, 9B ΔIDR2, 9B ΔIDR3, 9B RG5, and 9B RG9 coding sequences were subcloned into the pET-28a(+) (Novagen) bacterial expression vector to produce N-terminally 6xHis-tagged proteins containing a thrombin cleavage site, using *BsaI*-generated overhangs compatible with *NheI* and *BamHI* restriction sites. As a negative control, GFP coding sequence was cloned into the pET-28a(+) vector, resulting in the 6xHis-Thr-GFP construct. The constructs were expressed in E. coli BL21 (DE3) by induction with IPTG (1mM). Next, we proceed with the purification of the His-tagged proteins using Nickel-NTA affinity protein (ThermoFisher) following the manufacturer’s recommendations.

### Protein extraction and Western Blot analysis

Western blot analyses were performed to monitor the accumulation of purified recombinant proteins from *E. coli* and to verify the *in vivo* expression and stability of the constructs in *N. benthamiana*. For *in planta* assays, total proteins were extracted from agroinfiltrated leaf discs homogenized in extraction buffer (1,5 M Tris-HCl pH 8.8, 4% (w/v) SDS, and 30% (v/v) glycerol) freshly supplemented with DTT. For bacterial samples, aliquots corresponding to the different purification steps were collected. All proteins samples were boiled in Laemmli buffer, resolved by 12% SDS-PAGE, and transferred to PVDF membranes (Amersham). Immunodetection was performed using specific primary antibody (anti-GFP), followed by incubation with corresponding secondary antibody. Signal detection was achieved using BCIP-NT substrate according to standard procedures.

### Northwestern Blot analysis

Northwestern analysis was carried out as previously described (Pallas et al., 1999). Purified 6xHis-Thr-GFP-9B WT, deletion mutants (6xHis-Thr-GFP-9B ΔIDR1, 6xHis-Thr-GFP-9B ΔIDR2, 6xHis-Thr-GFP-9B ΔIDR3), and point mutants (6xHis-Thr-GFP-9B RG5 and 6xHis-Thr-GFP-9B RG9) were separated by 12% SDS-PAGE and transferred to PVDF membranes. Membranes were incubated overnight at 4°C in renaturation NW buffer (10 mM Tris-HCl, pH 7.5; 1 mM EDTA; 0.1 M NaCl; 0.05% TRITON-X100; and 1% BSA). Next, the membranes were incubated with 20 mL of NW buffer (without BSA) containing DNase-treated AMV sgRNA 4 (1.5 µg) labeled with digoxigenin (DIG RNA labeling Mix, Roche, Mannheim, Germany) for 2 h at room temperature, followed by three washes with NW buffer containing 0.3% Tween 20. Then, the membranes were incubated for 1 h in 20 mL of NW buffer containing anti-DIG (1:10000) (Anti-digoxigenin-AP, Fragments Fab; Roche, Mannheim, Germany), followed by three washes. The membranes were revealed using BCIP/NBT colorimetric substrate (Thermo Fisher Scientific, Waltham, MA, USA) following the manufacturer’s guidelines.

### Confocal laser scanning microscopy and image analysis

For *in vitro* phase separation imaging, purified proteins (stored in lysis buffer containing 10 mM Tris-HCl pH 7.5 and 20 mM imidazole) were diluted to the final concentration (10 µM for standard assays, and 1-24 µM for concentration-gradient experiments). These dilutions, along with the respective chemical disruptors or crowding agents (1,6-hexanediol, PEG, or NaCl), were prepared in PCR tubes. A 10 µL aliquot of each mixture was immediately deposited onto a glass slide and sealed with a coverslip. For *in vivo* imaging, 1-cm diameter leaf discs were excised from agroinfiltrated *N. benthamiana* leaves at 48 h (for PEG experiments) or 72 h (for 1,6-hexanediol experiments) post-infiltration (hpi) and mounted in water on glass slides. To evaluate the role of RNA in ALKBH9B phase separation dynamics, varying amounts of sgRNA4 AMV (0, 500, and 1000 ng) were added to the purified protein mixtures prior to microscopic observation.

Images were acquired using either a Stellaris 8 FALCON (Leica) or an AxioObserver 780 (Zeiss) confocal microscope equipped with a 40x water-immersion objective. GFP fluorescence was excited at 488 nm, and emission was collected between 490 and 527 nm. Chloroplast autofluorescence was concurrently recorded (excitation 488, emission 674-715 nm). To accurately capture the intracellular distribution of the condensates, *in vivo* images were acquired complementary as Z-stacks and maximum intensity projections are displayed.

Quantitative analysis of the *in vitro* condensates was performed using ImageJ/Fiji software. Images were converted to 8-bit format and segmented by thresholding. Particle analysis was restricted to puncta with a size range of 0 to 30 µm^2^ and a circularity index of 0.5 to 1.0. The mean condensate area and spatial density (number of puncta per 10.000 µm^2^) were calculated. Statistical comparisons across different categories and treatments were performed using a one-way analysis of variance (ANOVA).

### Fluorescence Recovery After Photobleaching (FRAP) analysis

FRAP experiments were performed on *N. benthamiana* leaf discs using the AxioObserver 780 (Zeiss) confocal microscope with a 40x water-immersion objective. The FRAP module was configured to acquire three pre-bleach images. Subsequently, specific regions of interest (ROIs) corresponding to GFP:9B condensates were photobleached using 100 iterations at 100% laser intensity at 488 nm. Fluorescence recovery was continuously monitored at 0.3-0.4 second intervals for a total of 30 s post-bleaching using minimal laser power. For data processing, background fluorescence measured from a cell-free ROI was subtracted from the recovery values. Additionally, the fluorescence intensity of a nearby unbleached condensate was recorded over time as a control to correct for incidental photobleaching during the recovery phase. Quantitative analysis of recovery kinetics, including the mobile fraction and half-time of recovery (t1/2), was performed using GraphPad Prism software by fitting the normalized fluorescence data to a single-exponential decay model.

## ACKNOWLEDGMENTS

The authors thank L. Corachan-Valencia for technical assistance. This work was funded by grant PID2023-149245NB-I00 from the Spanish MCIU/AEI/10.13039/501100011033 granting agency and European Regional Development Fund (ERDF/10.13039/501100008530). V. G. is the recipient of a PhD fellowship from the Ministerio de Ciencia, Innovación y Universidades of Spain.

## AUTHOR CONTRIBUTIONS

VG-S, M-OL, and JAN: performed experiments; VG-S, M-OL, JAN and JAS-N, formal analysis; VP and FA: supervised the study; ALL, writing - review & editing; VP, FA and JAS-N, funding acquisition.

## DATA AVAILABILITY STATEMENT

All relevant data supporting the findings of this study are contained within the manuscript and its supporting materials.

## SUPPORTING INFORMATION

Additional Supporting Information may be found in the online version of this article.

## FIGURE LEGENDS

**Table S1. List of oligonucleotides used in this work.**

**Supplementary Figure 1. Amino acid sequence alignment of ALKBH9B WT and its mutant variants.** The sequences of ALKBH9B WT, ΔIDR1, ΔIDR2, ΔIDR3, RG9, and RG5 were aligned using Clustal Omega and visualized with ESPript 3.2. Strictly conserved residues across all sequences are highlighted with a red background, whereas non-conserved regions are shown with a white background. Deletions within the ΔIDR variants are represented by black dots. Alanine substitutions in the RG9 and RG5 mutants are indicated by black font color instead of red.

**Supplementary Figure 2. Purification of recombinant ALKBH9B protein variants.** Coomassie Brilliant Blue staining of purified recombinant proteins expressed in *Escherichia coli*. Lanes from left to right: 6His-Thr-GFP:HA, 6His-Thr-GFP:9B WT, and the mutant variants ΔIDR1, ΔIDR2, ΔIDR3, RG9, and RG5. Black arrowheads indicate the expected electrophoretic mobility of each purified protein.

**Supplementary Figure 3. Transient *in vivo* expression of ALKBH9B variants.** Western blot analysis confirming the expression of the recombinant proteins used for *in vivo* assays. Total proteins were extracted from *N. benthamiana* leaves transiently expressing the different variants. The upper panel shows the immunodetection using an anti-GFP antibody. The lower panel shows Coomassie Brilliant Blue staining as a loading control. Lanes from left to right: GFP:9B WT, the mutant variants ΔIDR1, ΔIDR2, ΔIDR3, RG9, RG5, and a negative control.

**Supplementary Figure 4. Detailed *in vivo* subcellular localization of ALKBH9B variants.** Representative confocal microscopy images of *N. benthamiana* leaf epidermal cells transiently expressing free GFP, GFP:9B WT, or the mutant variants ΔIDR1, ΔIDR2, ΔIDR3, RG9, and RG5. For each construct, the upper horizontal panels display GFP fluorescence and bright-field (BF) images captured at lower zoom, whereas the lower horizontal panels display images captured at higher zoom. Scale bars = 20 µm.

**Supplementary Figure 5. Subcellular localization of ALKBH9B driven by its native promoter.** Representative confocal microscopy images of *N. benthamiana* leaf epidermal cells transiently expressing GFP:9B WT or the GFP:9B RG5 mutant under the control of either the constitutive 35S promoter (p35S) or the 2.5 kb native *ALKBH9B* promoter (*pALKBH9B*). The upper horizontal panels show GFP fluorescence, and the lower horizontal panels show bright-field (BF) images. Scale bars = 20 µm.

**Supplementary Figure 6. Quantification of AMV RNA accumulation across independent biological replicates. (A-C)** Densitometric analysis of viral RNA levels from three independent northern blot bioassays performed in *N. benthamiana* leaves transiently expressing GFP:9B WT, the mutant variants ΔIDR3 and RG5, or an empty vector control. Panels (A), (B), and (C) represent individual bioassays with *n* = 5, *n* = 4, and *n* = 4 biological samples per condition, respectively. Individual data points represent independent biological replicates. Horizontal lines and error bars indicate the mean ± SEM. Statistical significance relative to the empty vector control was determined using one-way ANOVA followed by Dunnett’s multiple comparisons test (ns, not significant; \**p* < 0.05; \*\*\**p* < 0.001; \*\*\*\**p* < 0.0001).

